# Better and faster collective decisions by larger fish shoals in the wild

**DOI:** 10.1101/2024.10.14.618136

**Authors:** Korbinian Pacher, David Bierbach, Yunus Sevinchan, Carla Vollmoeller, Alejandro Juarez-Lopez, Jesús Emmanuel Jiménez-Jiménez, Stefan Krause, Max Wolf, Pawel Romanczuk, Lenin Arias-Rodríguez, Jens Krause

## Abstract

Studies on collective cognition have provided many examples of decision-making benefits in terms of animals sharing information about predators, prey or resources in their environment. It has been shown how the efficient spread of adaptive information within groups can provide benefits which increase with group size. Little is known, however, to which extent groups also amplify maladaptive information such as false alarms and whether such costs reduce or even nullify the above benefits. Here, we investigated fish shoals in the wild that responded collectively with escape dives when attacked by birds. We analysed the response of shoals in reaction to hard-to-detect bird attacks and similar but harmless flybys as a function of shoal size. With increasing shoal size fish increasingly detected predator attacks (true positives) while their false alarms remained constant. Therefore, larger shoals became better at correctly classifying potentially dangerous stimuli rather than becoming more sensitive to all stimuli potentially related to attacks. In addition, decision time decreased with increasing shoal size. Larger shoals were thus able to mitigate two major trade-offs inherent in solitary decision making: the trade-off between true and false positives and the trade-off between speed and accuracy. We report performance increases at shoal sizes of tens of thousands of fish and pose challenges for the modelling of the underlying mechanisms.

## Introduction

Free living animals are under constant pressure to make decisions in which errors can have serious consequences, most crucially when assessing the presence or absence of imminent predation risks ^1^. In animals groups, decision quality can be improved through emergent collective cognition (e.g., the integration of independently collected information through social interactions resulting in an adaptive group-level decision-making processes), a process observed in many social vertebrate taxa including humans ^2–7^. With increasing group size collectives cannot only increase the probability of correct decisions, but also reduce decision time ^8^. However, most of the supporting evidence was obtained from theoretical considerations or laboratory studies ^6,7,9,10^. Yet it is unclear if decision accuracy (quality) and speed indeed increase with group size under field conditions when noisy environments introduce misleading information that can hamper correct assessments of cues (an aspect often unaccounted for in laboratory settings like applied in ^8^, see also: ^11,12^).

In detection theory the assessment of a binary choice situation can result in two errors: A false positive (*false alarm*) in which a condition (i.e., predation risk) is absent, but a response (i.e., anti-predator behaviour) was wrongly initiated and a false negative (*miss*) in which a condition is present, but no response was initiated ^13^. In this framework a correct decision is either a true positive (*hit*: condition and response present) or a true negative (*correct rejection*: condition and response absent). A solitary animal can increase its general cue sensitivity to increase true positive probabilities ^14,15^. However this raises the probability for false positives, not improving overall accuracy ^7,16^. To really improve the accuracy of decision-making means to manage the trade-off between true and false positives effectively and decouple the probabilities of both response types.

A common strategy to achieve higher decision accuracy is by accumulating more and potentially also better information from the environment ^17^. Yet longer sampling increases the second crucial decision parameter: Speed. A higher decision quality is therefore often associated with a longer decision time, a relationship referred to as ‘speed-accuracy trade-off’^18^. To explore if a decision by a group is indeed more accurate and faster requires information on both trade-offs. Thus, in a first step accuracy must be measured in the form of true *and* false positive probabilities (and their negative counterparts respectively). This is a prerequisite so that measurements of decision time can provide information on how the speed-accuracy trade-off is managed. There has been some (mostly theoretical) exploration of how collective cognition might mitigate the general accuracy trade-off ^2,3,7,9,19–22^. However, predictions are less clear for the speed accuracy trade-off ^11,23–25^. Empirical evidence exploring a combination of both trade-offs in relation to group size is very limited and practically non-existent for animal groups under natural conditions.

A key difficulty in observing true and false positives in the wild is the limited knowledge of the environmental state—for example, whether a predator is present and whether it poses enough danger to justify an anti-predator response ^26^. Although this may seem straightforward, most field studies rely on quantifying prey responses rather than actual environmental states to assess true and false positives (e.g., a fleeing prey group is recorded, and the situation is *then* classified based on whether the scientific observer and not the prey could detect a predator). This approach usually fails to account for negative responses, such as correctly ignored harmless stimuli (true negatives) or overlooked predators (false negatives), which remain unrecorded as they do not result in visible prey responses. However, these negatives are crucial for determining whether a stimulus was assessed correctly and for a calculation of overall decision accuracy. A second issue is that most observational studies cannot account for the total number of decision-triggering events during the observation period, which is necessary for calculating the reliable probabilities for decision outcomes. Finally, it is challenging to determine the time in which a decision was made without knowing when and if the prey group perceived a stimulus.

Our study system is suitable to address these issues. We explore collective decision-making in large shoals of sulphur mollies (*Poecilia sulphuraria*) - small livebearing fish - inhabiting sulphidic springs in the South of Mexico which are under regular predation by birds. In their extreme habitat, H_2_S-induced hypoxia especially during the day leads to the formation of large surface bound fish shoals since fish skim the oxygen-rich water surface through aquatic surface respiration ^27–29^. These shoals attract a variety of predatory bird species, which usually prey on shoals in consecutive sequences of attacks (henceforth called ‘attack sequence’, ^30,31^). To counter this frequent type of predation, sulphur mollies have evolved a unique collective behaviour: When disturbed, fish collectively dive down in a staggered fashion, creating ripples at the water surface that appear as quantifiable surface waves. After an initial collective dive, fish shoals often repeat this behaviour for several minutes, creating repeated surface waves (from here on referred to as repeat waves), a phenomenon that has been shown to decrease a predator’s attack frequency and success and can even lead to a predator abandoning its target shoal ^30^. In addition to high rates of bird predation, sulphur mollies are further confronted with harmless disturbances like moving tree branches, reed grass or flybys of non-attacking birds for which collective (repeated) diving would mean a waste of energy, and therefore a wrong response ^32^. While an initial reaction dive is reported for both disturbances, repeat waves were previously associated mainly with real attacks on the shoal ^30^. So, the decision to continue with anti-predator behaviour is somewhat independent of the initial diving response that can be seen as indication of a disturbance detection. As most bird attacks are conspicuous - the attacking predators usually enter the water with large parts of their body (e.g., kingfishers) and are therefore easily detected ^32^ - the decisions to repeatedly dive or not are relatively straight forward. However, the great kiskadee’s (*Pitangus sulphuratus*) attack style in which the bird attacks in full flight and only its beak briefly penetrates the water surface, adds ambiguity ^30^. This attack closely resembles a harmless flyby, and it is expected to present the prey fish with discriminatory challenges, operating in a cue parameter space in which true and false positives are both possible. By focusing on this predator, we investigated if larger shoals could increase their true positive rates and decrease false positive rates in the form of discriminating more accurately between a real attack and a harmless flyby. Additionally, we examined if any accuracy gains increased the decision time or if larger groups could make both better and faster decisions. In knowing the true environmental state, namely if a disturbance represented a kiskadee attack or a harmless flyby (Fig. 1), we could apply a detection theory framework and classify discrete decisions by the prey to proceed with collective waving as correct or false (Fig.1 d, e). Additionally, we measured decision speed as the time interval between initial detection (reaction dive) and the start of the repeat waves (Fig. 1c). This enabled us to collect data on both relevant decision parameters: Speed and accuracy. Lastly variations in shoal sizes provided an opportunity to investigate the relationship between collective decision performance and group size.

**Figure 1:**
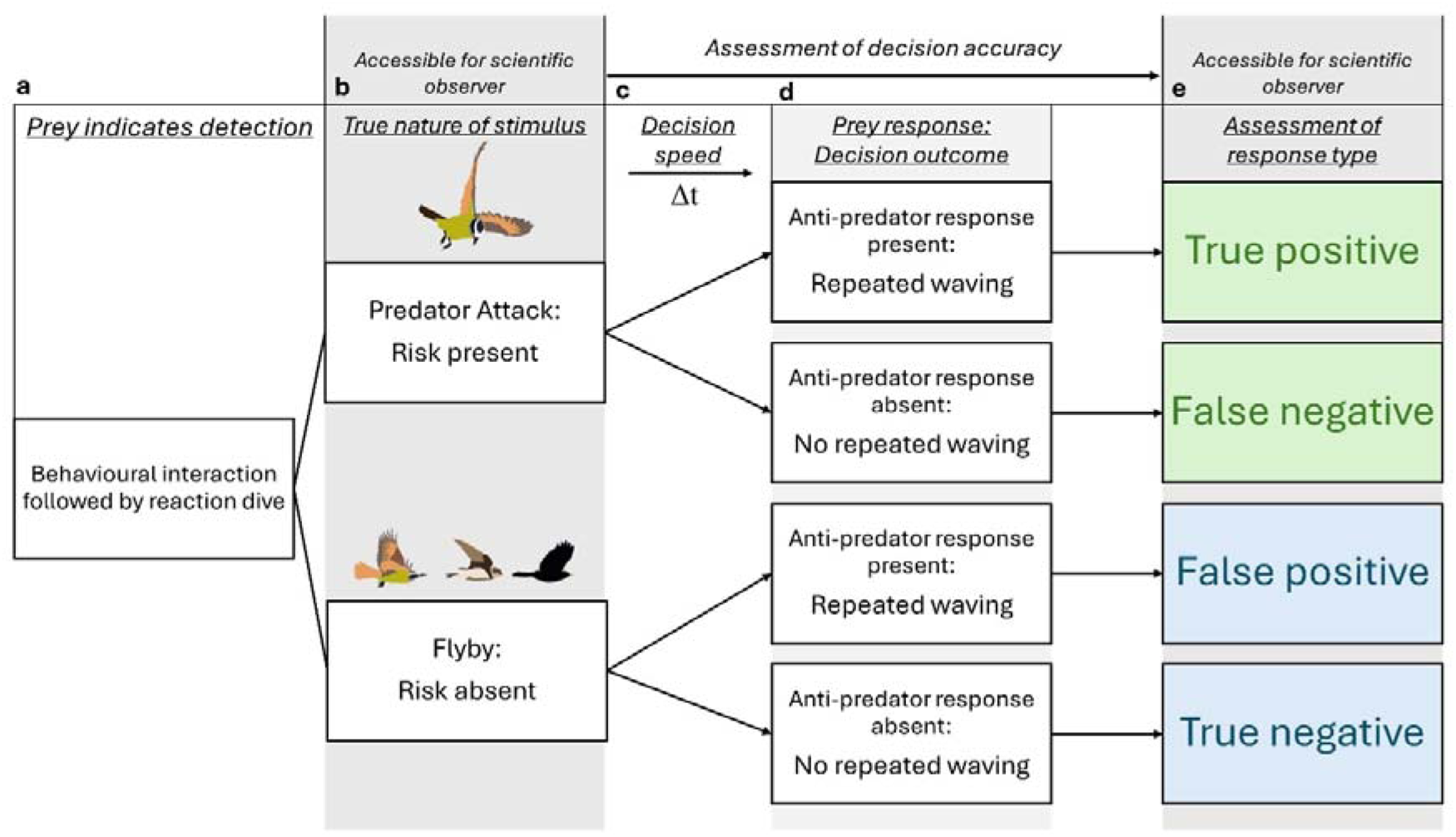
Key system features enabling a comprehensive approach to investigate collective decision accuracy and decision time under field conditions. a) A reaction dive by the prey indicates that a stimulus was detected and represents the start of a discrete situation requiring a decision by the prey. b) While overflight attacks and flybys present prey groups with discriminatory challenges, their true nature is clear to the scientific observer, which is a prerequisite for the classification of a decision. c) The time interval between the initial reaction dive and the start of collective anti-predator behaviour represents a direct measurement of decision time. d) The prey group indicates through the presence or absence of repeated waving if a stimulus was assessed as dangerous or harmless. e) In knowing the true nature of a stimulus in combination with the prey’s reaction a scientific observer can accurately assess all four possible response types. This crucially does not only include positives in which a reaction is shown but also negative responses.

## Results

### Variation in group sizes and fish density (shoal size)

As sulphur molly shoals consist of a single layer of fish (See inlay in Fig. 2a), we used shoal size area as a proxy for shoal size because of the huge numbers of fish involved making direct counting impossible. Measured shoal size area ranged between 31.2 m^2^ and 75.03 m^2^ (See Fig. 2 e) and mean nearest neighbour distance was at 16.29 ± 3.76 mm (N = 64508 fish positions in 76 frames analysed). We found significant differences in fish nearest neighbour distances among our study sites but no relationship between shoal size area and fish density (N = 76; GLMM_gaussian_ shoal size area: χ^2^(1) = 1.0, p = 0.30; location: χ^2^(2) = 69.1, p < 0.001; See Fig. S4 in supplementary material). The swimming direction (polarisation) of the fish in the shoals was neither influenced by shoal size area nor side location (N = 76; GLMM_gaussian_ shoal size area: χ^2^(1) = 0.09, p = 0.75; location: χ^2^(2) = 2.81, p = 0.24). However, a marginal significant interaction-term indicates that the effect of area on mean polarization may differ depending on the site (N = 76; GLMM_gaussian_ shoal size area*site χ^2^(2) = 5.99, p = 0.049).

**Figure 2:**
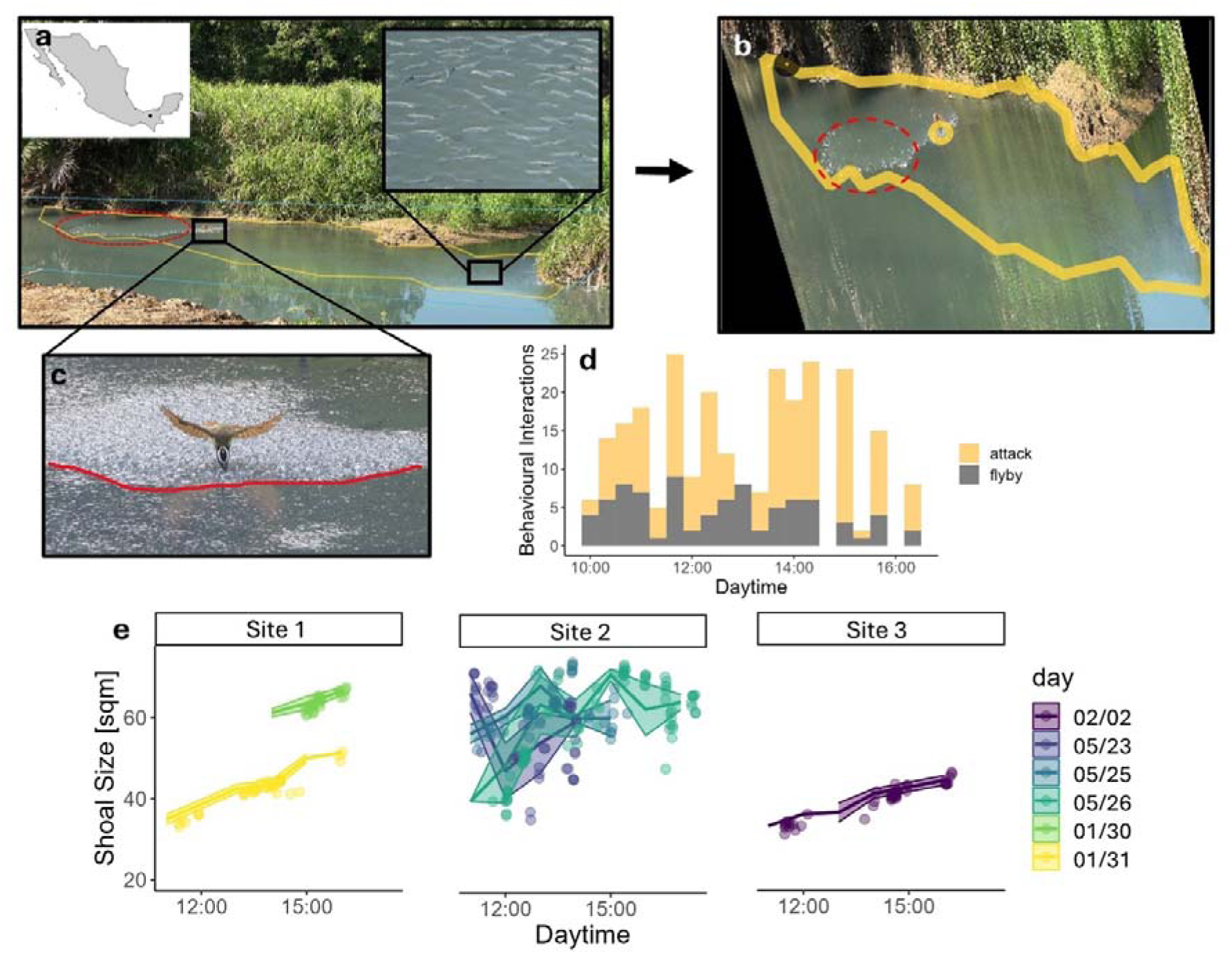
Study system overview. (a) Field work was conducted along the ‘El Azufre’ sulphidic riverine complex in Tabasco, Mexico, where sulphur mollies forming large shoals at the surface can be video recorded from the riverbanks as depicted in a still frame from a typical recording, with the shoal outline marked in yellow. (b) Spatial measurements like shoal size area are extracted from rectified frames of video recordings; here, the same still frame as shown in (a) is depicted after perspective transformation. (c) Great kiskadee during a typical overflight attack. Note how only the bird’s beak penetrates the water surface and how fish show an initial diving response during the attack. Diving fish can clearly be distinguished by the increased reflection following water disturbances as marked in red in a, b, c. (d) Bird attacks and flybys over the course of the day (all sampling days combined). (e) Shoal size measurements for the three sampling locations. Lines indicate hourly mean shoal size per day with shaded area as standard deviation. Dots represent raw data (N = 258, see supplementary Table S1 in for more information on sampling locations).

### Decision accuracy and group size: True and false positive rate

We analysed decisions by fish shoals in response to 177 kiskadee attacks and 81 flybys. Sulphur molly shoals in general showed a significant increase in their true positive rate with increasing shoal size for kiskadee attacks, with some variation of effect strength among shoal locations (N = 177; GLMM_binomial_ shoal size area: χ^2^(1) = 18.14, p < 0.001; Fig. 3a in green; location χ^2^(2) = 18.4, p < 0.001; Fig. 3c, no effect of attack in sequence: χ^2^(1) = 0.32, p = 0.5). In contrast, sulphur molly shoals’ false positive rates were not influenced by shoal size area, location or flyby bird species (N = 81; GLMM_binomial_ shoal size area: χ^2^(1) = 0.03, p = 0.86, Fig. 3a in blue; location χ^2^(2) = 0.58, p = 0.74; species: χ^2^(1) = 0.97, p = 0.32). Given that we had a larger sample size testing for true positive rates compared to false positives, we performed a bootstrapping simulation (see method section for details). Our results suggest that the non-significant result of shoal size for false positives is likely due to the true absence or a very small magnitude of the shoal size effect, rather than being a consequence of the smaller sample size (See supplementary figure S3). In contrast to kiskadee attacks, kingfisher attacks were always followed by repeat waves and thus classified as true positives (See Fig. 3b), confirming our prediction that the detectability of kiskadee attacks ranges between false positive rates for flybys and true positive rates for kingfishers (see introduction).

**Figure 3:**
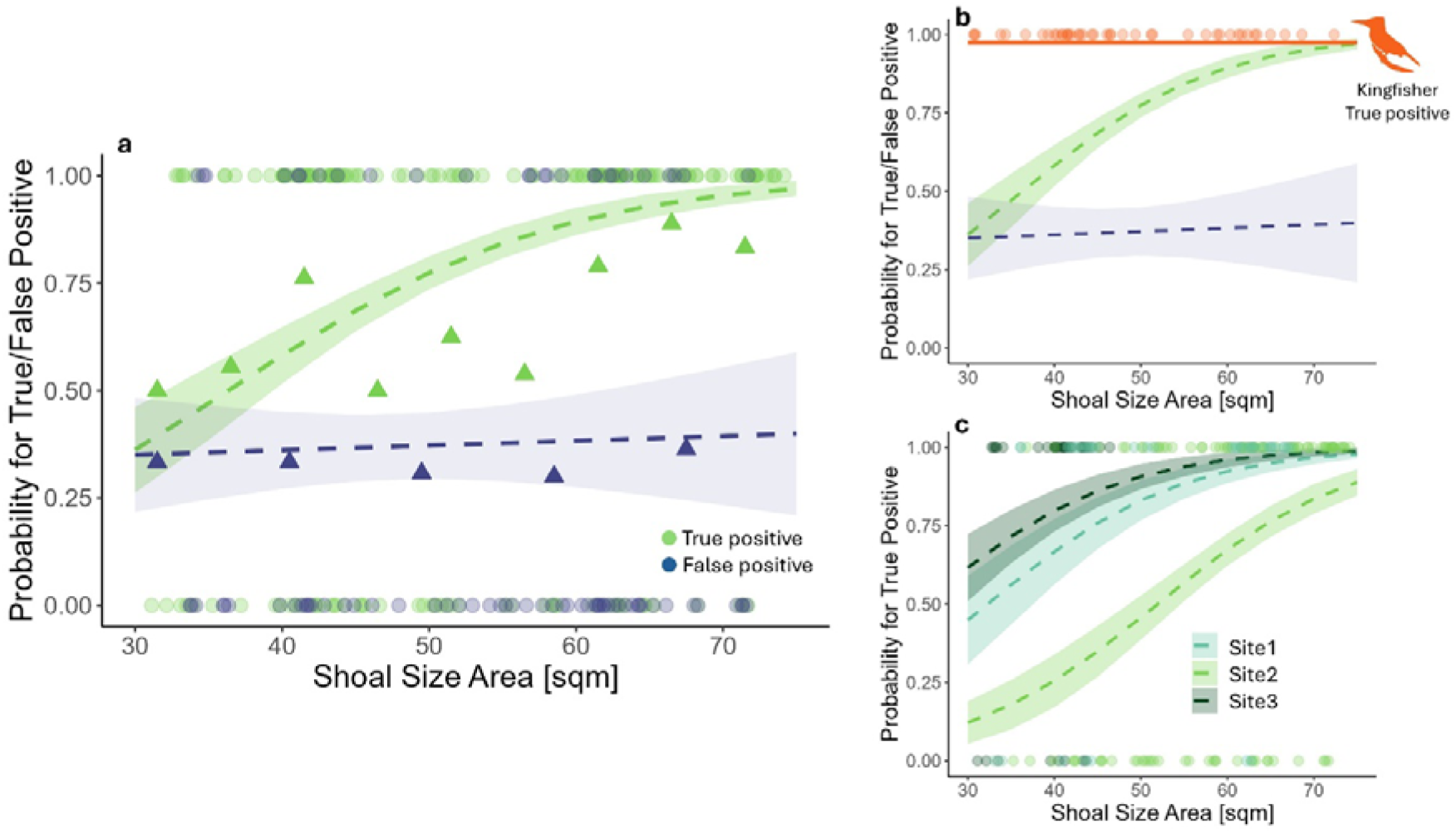
Effect of shoal size area on decision accuracy. True and false positive probabilities during evaluation of bird attacks by sulphur molly shoals (a) When attacked by kiskadees, the true positive probability (= correct repeated diving response; N = 177 cases; green) significantly increased with size of the shoal while the false positive probability (=wrong repeated diving response to a flyby; N = 81, blue) remained unaffected. Triangles represent binned mean true/false positive rates. (b) Comparison of the effect of shoal size on true positive probability (green) for kiskadee attacks and kingfisher attacks (N = 47, orange) as control. Even small shoals always showed repeat waves after conspicuous kingfisher attacks (full body plunge-in attacks), leading to a perfect classification. False positive rate for flybys (blue) is included for comparison, note how true positive rate for ambiguous kiskadee attacks changes in between both constant probabilities. (c) Differences of the shoal size effect on the true positive probability for kiskadee attacks by sampling location. Dots represent raw data, and dashed lines represent model predicted regression fits with SE on raw data.

True positive responses were characterized by a significantly higher wave number (mean TP wave number = 5.65 ± 5.16) than found for false positives (mean FP wave number = 2.04 ± 1.53), an effect that scaled up with shoal size area and differed among locations (N = 153; GLMM_negbinom_, shoal size area: χ^2^ (1) = 8.04, p < 0.004; response type: χ^2^ (1) = 13.60, p < 0.001 see Fig. 4b in green; location χ^2^(2) = 7.43, p = 0.024). When true and false positives were investigated separately, the influence of shoal size area and location on repeat wave numbers was only significant for true positives (N = 127; GLMM_negbinom_ shoal size area: χ^2^(1) = 8.63, p = 0.003 see Fig. 4a in green; location χ^2^(2) = 7.9, p = 0.019), while the wave numbers for false positives were not influenced by shoal size area or location (N = 26; GLMM_negbinom_ shoal size area: χ^2^(1) = 0.16, p = 0.69; location χ^2^(2) = 0.75, p = 0.68; see Fig. 4a in blue). With increasing shoal size, the time between the end of the initial detection dive to the first repeat wave decreased, meaning that larger shoals decided faster to proceed with anti-predator behaviour. This was independent of response type and location. (N = 148; GLMM_gaussian_, shoal size area: χ^2^ (1) = 9.77, p = 0.002; location: χ^2^(2) = 5.50, p < 0.063 response type: χ^2^ (1) = 1.29, p = 0.25 see Fig. 4c, d). Predator success and waiting time were not influenced by any of our predictors such as the number of waves a great kiskadee had experienced before its attack, shoal size area, decision time or location (see supplementary information for model details: M7 and M8).

**Figure 4:**
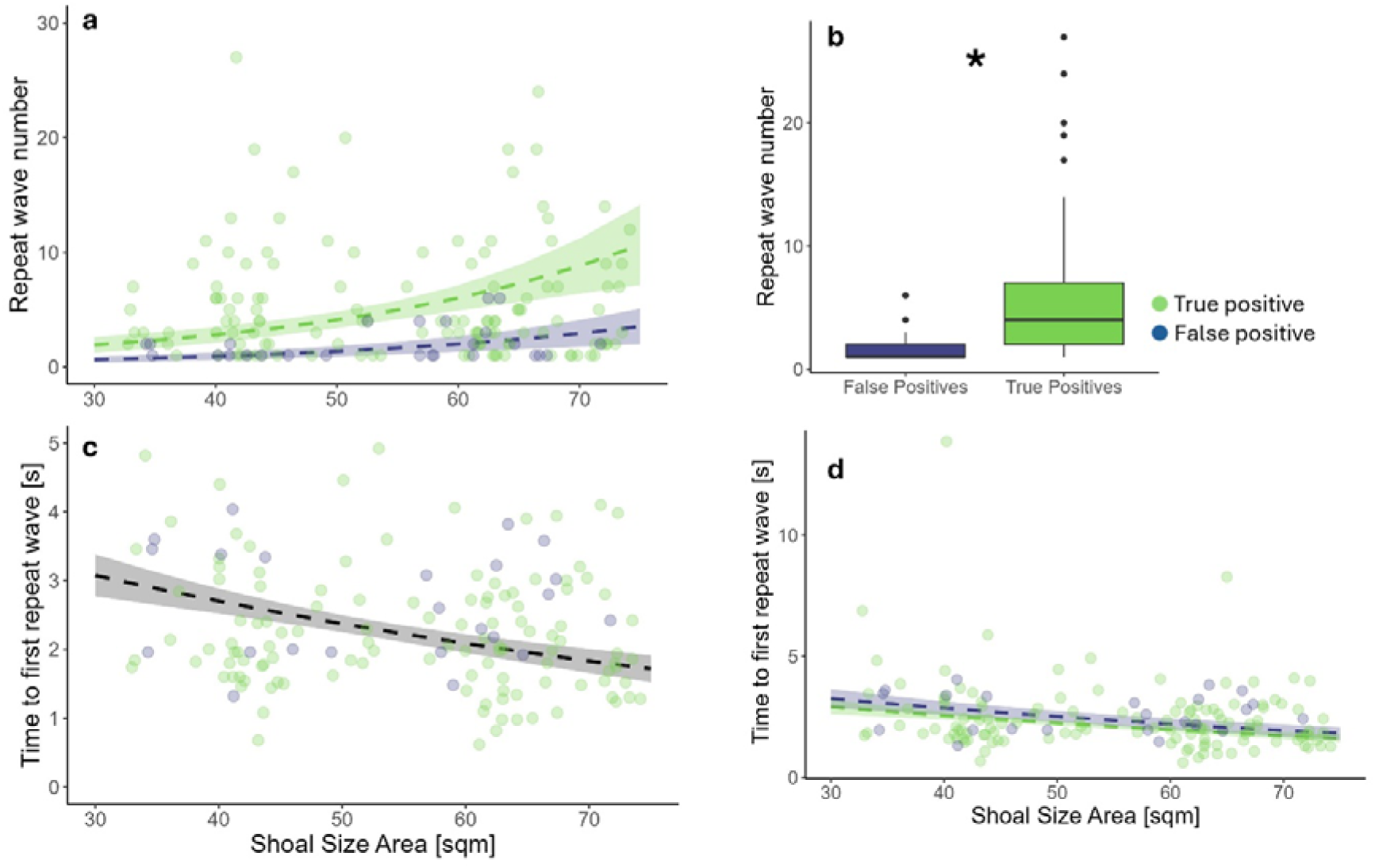
Effect of shoal size area on the magnitude of collective waving and decision time. (a) With increasing shoal size area wave numbers increased for true positive responses but not for false positives (green: true positive, N = 127 / blue: false positive, N = 26). Response type and area had a significant effect on the number of repeat waves. (b) When within a true positive sulphur mollies showed significantly more repeat waves compared to a false positive reaction. (c) With increasing shoal size decision time decreased (N = 148) expressed as time to first repeat wave. Decision time significantly decreased with shoal size, but (d) was not affected by response type (green: true positive / blue: false positive). Dots in a,c,d represent raw data and dashed lines represent model predicted regression fits with SE on raw data, note the cropped y-axis for greater clarity of statistical effect in c. Boxes in b indicate the inter-quartile range (IQR), with the central line depicting the median and the whiskers extending to 1.5IQR. * denotes statistically significant effect.

## Discussion

Larger shoals discriminated more accurately between harmless bird flybys and dangerous attacks by a hard-to-detect predator than smaller shoals. As the size of fish shoals increased, so did probabilities for true positives. Importantly, this increase in correct decisions did not come with the cost of a higher probability for false positives, indicating a genuine enhancement in decision accuracy. Additionally, larger shoals demonstrated faster decision-making by initiating anti-predator responses quicker than smaller shoals. Our work therefore provides novel evidence in an ecologically relevant context in the wild that collectives can produce better and faster decisions.

Our results show that while true positive response probabilities increased—indicating better detection of kiskadee attacks—there was no corresponding rise in responses to harmless stimuli as shoal size increased. This suggests that larger shoals became better at categorising potentially dangerous stimuli rather than just becoming more sensitive to them. Kiskadee attacks closely resemble harmless flybys, as the birds only slightly dip their beaks into the water, making their attacks harder to distinguish from non-threatening disturbances compared to other predator attacks like those of kingfishers. This may explain why Doran et al. (2022) reported minimal collective waving responses to kiskadee attacks, assuming sulphur mollies always treated these attacks as harmless flybys. This is not the case as we show in our current study but likely went unnoticed by Doran et al. (2022) as their study focused on smaller shoals with minimal size differences. Our findings reveal that the largest observed shoals could detect kiskadee attacks with a level of accuracy similar to that observed for other predators, like kingfishers, whose attacks produce more intense sensory cues ^32^. Even small shoals nearly always detected kingfisher attacks, responding with ten times more waves on average compared to kiskadee attacks ^30^. This difference in response intensity may explain why we did not observe a negative impact on success or waiting time from increased wave numbers, somewhat contrasting to Doran et al. (2021), who found a strong negative effect on kiskadee hunting performance when wave numbers were experimentally elevated to levels typical of kingfisher attacks. We argue that sulphur mollies evolved their repeat wave behaviour in response to specialized piscivores like kingfishers, herons, and egrets, rather than opportunistic predators like kiskadees. Still, when attacked by kiskadees the sulphur mollies’ increased efficiency to detect threatening attacks with increasing group sizes becomes apparent, even if the associated stronger response has only a subtle impact on the kiskadee’s hunting performance. A larger sample size might help to detect potential negative effects of increased waving on the kiskadee hunting success.

Larger groups of sulphur mollies were able to mitigate two major trade-offs inherent to solitary decision making: The trade-off between true and false positives and the trade-off between speed and accuracy ^2,7,20^. Previous considerations identified quorum-based decision rules as a mechanistic explanation of how collectives achieve this self-organized improvement in decision performance. In a quorum decision an individual responds to social information only when a behaviour is observed in a threshold number of nearby group members ^9^. This basic mechanism of social information use has been shown to improve collective decision making in a variety of taxa from eusocial insects ^33^ up to complex and highly relevant decisions in humans ^3,10,34^. Notably the effects of quorum rules and increasing group sizes in predation contexts substantially overlap with our findings such as faster decisions ^8^, higher true positive rates ^6^, constant false positive rates ^35^, better discrimination ^5^ and a decoupling of true and false positives i.e., a higher overall decision accuracy ^7^. Mechanistically, with increasing group size it becomes easier to reach a critical quorum threshold as typically more members perceive the triggering stimulus and by combining both personal and public information a correct response is facilitated. Although we determined overall group size (as shoal area that correlates with number of individuals in the shoal), our current study did not determine the number of responding (diving) neighbours a sulphur molly needs to initiate a dive itself and we thus can only speculate whether the observed increased decision efficiency in larger shoals is really caused by quorum-based decision-making.

Most laboratory studies report near-optimal decision accuracy in groups of a few dozen individuals ^5,7,8^, where the individuals can access information of all members (all-to-all coupling). Therefore, Ioannou (2017) suggested that self-organized cognition can only be confirmed if performance improvements are consistent across much larger group size changes, which is rarely the case in laboratory settings (see ^37^ for an exception). In our study, shoals consist of tens of thousands if not hundreds of thousands of individuals spread over entire river sections, circumstances under which all-to-all coupling is highly unlikely. Studies on behavioural contagion further challenge the notion of quorums as binary thresholds that lead to behavioural adoption on a collective scale once it is reached ^38,39^. This is particularly relevant for large collectives where temporary density fluctuations and visual occlusion may affect information spread, factors often overlooked in conventional quorum models ^39,40^. And while alternative models have worked towards integrating traditional majority rules with a more realistic network topography, they cannot yet supply plausible mechanistic explanations of how collectives can increase their decision performance as observed in our study ^25,41^. Recent work suggests that sulphur molly shoals resemble a system operating under a regime of self-organized criticality, a state allowing biological collectives, like neuronal networks, to optimize information processing, widely associated with the emergence of collective cognition ^27,42^. Operating at a critical point can influence true and false positive rates and decision speed to varying degrees ^39^ but does not yet explain how these responses can be decoupled effectively. However, extending such models of collective information processing to include dynamic sensitivity thresholds—like those observed in free living coral reef fish, which can selectively reduce sensitivity to misinformation ^43^ — may offer a promising direction for future research. In this context, our results, while providing only limited mechanistic insights, highlight the importance of studies on large collectives under field conditions for eventually understanding alternative mechanisms of emergent collective cognition and the ecological factors that led to their evolution.

The low and stable false positive probabilities across group sizes contrasts sharply with existing literature, which often report probabilities for false positives well beyond those for true positives ^15,44,45^. It is widely assumed that adjusting a general sensitivity threshold is the only way for animal groups to optimize predator detection while keeping the speed of decision constant ^46^. By setting thresholds to detect nearly every disturbance, groups are expected to accept the costs of frequent false alarms under the “better safe than sorry” principle ^47^, where high true positives outweigh the costs of false positives. However, this also assumes that groups face the same trade-off as a solitary decision-maker and can increase their true positive rates only at the cost of heightened false alarms. Our results clearly show that this is not the case and animal groups in the wild can decouple both responses. And while we are aware of the pitfalls that might arise by basing a conclusion on the absence of a statistical effect (i.e., no change in false positives), we are confident that a similar steep increase in false positives would have not remained undetected. Moreover, Gray and Webster (2023) argue that the costs of false alarms are difficult to measure and likely vastly understudied. We agree and suggest that in many environments where false alarms are costly, prey groups face selection pressure to develop mechanisms that achieve both high predator detection and low false alarm rates. This appears to be the case in our system, where low oxygen concentrations make repeated diving costly. The phenomenon of collective repeat waves in sulphur mollies is a somewhat regional phenomenon, however collective waves and similar manoeuvres as anti-predator behaviour are well documented from other fish species ^48^, birds ^49,50^ and insects ^51^. We therefore advocate for more comprehensive approaches and novel study systems to further unravel how collective cognition might shape behaviour in different environments. These insights are crucial to inspire future models of collective behaviour exploring the possibility of unique mechanistic aspects emerging from very large collectives.

## Material and Methods

### Study system

Fieldwork was carried out at the El Azufre riverine system in Teapa, Tabasco, Mexico. Here, geothermally heated sulphur springs flow into a natural freshwater stream leading to a sulphidic river stretch inhabited only by two sulphur adapted fish species, the widemouth gambusia (*Gambusia eurysthoma*) and the sulphur molly *(Poecilia sulphuraria*). The sulphur molly is most common and we thus refer to the shoals of fish in our study as sulphur molly shoals (see: ^52^. Fieldwork was conducted in January 2023 and May 2023 which represent the local dry season. During this time, temperatures rise over the course of the day and increasingly hypoxic conditions in the sulphur-affected river stretches lead to the formation of large surface bound shoals of sulphur mollies (for an extensive description on environmental conditions see ^28^ for water parameters and ^29^ for thermal regime). As the fish perform aquatic surface respiration to skim the oxygen rich surface layer, sulphur molly shoals exist in a quasi-two dimensional state and can comprise hundreds of thousands of individuals covering whole river stretches of up to 100 m^2^ ^27–29,53^. These shoals form consistently on a daily base at the same locations and can therefore be observed in a standardized way. However, while the formation of shoals on certain locations is highly predictable their size and densities can vary substantially over the course of the day and in between days ^52^.

Surface bound shoals attract a variety of avian predators such as kingfishers, kiskadees and herons ^31^. As hypoxia always forces fish back to the water surface these prey shoals are constantly exposed to a high predation risk. Thus, avian predators often approach and select a focal shoal which is then attacked multiple times in quick succession before the predator leaves ^28,30^. In addition to predator attacks, sulphur mollies are constantly confronted with randomly occurring harmless stimuli ^32^. A typical example is a bird flyby, which may represent a disturbance to the fish, but with a low likelihood to be directly followed by an attack as sulphur molly shoals are constantly exposed to flybys by traveling harmless and predatory birds alike ^32^.

The great kiskadee presents an ideal case as a predator. Unlike specialized piscivorous birds, it conducts a unique overflight attack, where only its beak briefly penetrates the water surface. This attack lacks typical predatory cues observed in more specialized predators ^32^, such as the slow approach by larger herons or the vigorous plunge diving by kingfishers (^30^, see also supplementary Fig. S2). The attack of the kiskadee resembles harmless flybys of other birds (which frequently take place) and therefore presents fish shoals with a high level of uncertainty that can be utilized to investigate the classification ability of fish shoals as a function of shoal size.

### Recording of predator-prey interactions in the field

To observe predator-prey encounters between sulphur mollies and the great kiskadee, we filmed a focal fish shoal in their natural habitat. To capture predator-prey interactions and size variations on the same focal fish shoals, we filmed at three different locations continuously with a fixed camera setup (25 FPS, Sony HD Handycam FDR-AX33) for complete days from 9:30 PM to 4:30 PM. During this time predators would arrive at and attack the prey shoals at will, which was then recorded on video. In addition, the occurrence of all attacks their outcome were recorded by a human observer.

### Video processing and analysis

As videos of predator-prey interactions are recorded from the shoreline, a first processing step was to correct for the oblique angle. To that end, a quadratic reference object of known size was placed onto the water surface at the end of every recording session– this enables a perspective transformation (“rectification”) using the OpenCV Python Library (Computer Vision Library; https://opencv.org/), augmenting a top-down view onto the shoal, where distances on the water surfaces can be measured in physical units (For full documentation see^54^).

Sizes of sulphur molly shoals were measured as water surface area ‘covered’ by fish. This is possible as the absence of oxygen in the system forces fish to perform aquatic surface respiration which leads to large shoals consisting of only a single layer of fish; subsequently, the spatial extent at the water surface can reliably serve as a measure of group size. This shoal size area is measured by manually defining a polygon that traces the outline of the shoal in frames of the original video; these points in the unrectified frames where then transformed into coordinates in the top-down view using the same perspective transformation procedure. Shoal outlines can be detected in video recordings by surface ripples originating from individual fish performing aquatic surface respiration. In addition, when shoals perform repeat waves, the extension of waving serves as an indication to reliably define shoal outlines (See Fig 2a, b). To account for measurement error, ten frames showing shoals of different sizes were measured five times each in random order. This revealed a mean measurement deviation of ± 2.35 m^2^ which represents a range between 3 – 7 % of total shoal size.

In the recorded videos, we noted to the frame each kiskadee attack or bird flyby (*behavioural interaction type*) using a custom video annotation software ^54^. An attack was defined as any interaction between a predator and the focal prey shoal in which the predator penetrated the water surface with its beak to grab a fish. A flyby was defined as any interaction between bird and the focal prey shoal without contact of the bird’s beak and the water surface and therefore no imminent predation risk for fish. Flybys could therefore be any case in which predatory or harmless birds passed through the area without attacking. We further scored the *number of collective waves by the prey* (wave number), the *size of the prey shoal* (area in m^2^) at the time of event and the *decision time* (seconds) as time interval from the end of the initial reaction dive to the first repeat wave. For attacks we recorded four additional variables: Firstly, a binary variable *success* (yes/no), which indicates, whether the attack was successful. Secondly, we attributed attacks to individual sequences of subsequent attacks by the kiskadee. For these sequences we calculated *inter-attack waiting time* (time between two successive attacks in seconds). A sequence was initiated with a first attack and ended when the bird left the location or waited more than 300 seconds before it would attack again, which represents the maximum observed inter-attack waiting time in previous studies ^30^. Introducing this time threshold was necessary as birds would not always leave after an attack sequence but stay in the area and rest or preen.

For every attack we recorded a *unique sequence id* (sequence_id) and the *attack number within its unique sequence* (number attack in sequence). For flybys we recorded *species* (kiskadee / other) as an additional variable. Lastly, we scored the prey reaction as a true or false positive or the corresponding negatives respectively (see Fig. 1).

For the investigation of decision accuracy, we focused on attacks by the great kiskadee as this predatoŕs attacks are expected to be highly ambiguous. A *true positive* was defined as repeated waving after an attack while a *false positive* was defined as repeated waving after a flyby.

We screened field recordings of eleven days comprising of approximately 70 hours of video footage. Repeated predator-prey interactions between the great kiskadee and shoals of sulphur mollies could be observed on a total number of six days (3 days in February 2023 and 3 days in May). Overall, we recorded 324 behavioural interactions (193 predator attacks and 84 flybys; 47 kingfisher attacks) equally spread over the whole day (See Fig. 2d). To test our prediction, that kiskadee attacks are special in the regard that they present prey shoals with an unusual high level of uncertainty compared to other predators we randomly selected 47 attacks by kingfishers as a control group (See supplementary information for more information). According to our predictions and observations from Doran et al. (2022) the characteristic plunge diving attacks by kingfishers should be correctly detected even by relatively small shoals and therefore have a high and constant true positive rate. To rule out that the prey behaviour in reaction to kiskadees or overflights was influenced by other disturbances (e.g., herons or other wildlife) we removed all kiskadee interactions that were preceded or succeeded by another disturbance within a 60 second time window (16 kiskadee attacks and 3 flybys). When this was the case the present attack sequence was treated as if it had ended and the next attack by a great kiskadee started a new sequence. To avoid any observer bias, the number of repeat waves and the shoal size were measured in a double-blind procedure and in random order.

### Fish density measurements

We further measured fish density from close-up video recordings for all our field sites. These recordings were done on the same focal shoals as the ones at which predator-prey interactions were filmed, but on different days. This was necessary to avoid attacking birds being disturbed by the camera setup which had to be placed near the focal shoals to allow close-up recordings. We recorded focal shoals on two days per location at every hour from 10:00 AM to 4:00 PM for sessions of ten minutes. These videos were then rectified as described above and two frames per session were randomly selected. We used a custom semi-manual computer vision processing pipeline to mark every individual fish, its position and orientation within each frame (published in ^52^). Based on the annotated positions and orientations, we calculated the nearest neighbor distance (NND - center of head to center of head) and the relative swimming polarization (tail to head centered unit vector) in every frame. Shoals used for detailed density recordings were additionally filmed with a wider camera angle that allowed for the quantification of shoal size area to relate changes in density and polarization to the area covered by fish.

### Statistical analysis

All statistical analysis was performed in R (v.4.1.2). We fitted general linear mixed models (GLMM) with the glmmTMB library ^55^. In the main text we report the Wald χ2 statistics (two-sided) for assessment of the significance of fixed effects in final constructed models, while detailed summaries of final models can be found in the supplementary material. Unless specified otherwise all models were fitted *with shoal size area* and *location* as fixed and sampling day as random effect. *Sampling location* was included as fixed effect, as this variable had only three levels and was therefore not suited to be included as random intercept. Sampling day was included with a random effect structure nested in location, as every sampling day was filmed at one location only.

### Group size

As fish densities and polarization were measured on separate days, we first investigated the relationship of *density* and *polarization* with *shoal size area*. Because of a strong correlation between density and polarisation both variables were examined in separate models with a gaussian error distribution (identity link).

### Decision accuracy

To test for the influence of shoal size on the true positive rate, *true positive* (true positive: yes / false negative: no) was fitted as response variable in a logistic regression with logit link function. In addition to *shoal size area*, *attack number in sequence* was included as fixed effect. To test for the relationship of shoal size and *false positive* rate we fitted a logistic regression (link = logit) including *false positive* (false positive: yes / true negative: no) as our binary response variable. In addition to *shoal size area* we included *flyby species* as fixed effect to account for differences in false positive rates between flybys by the kiskadee and other birds. As our true positive data set contains substantially more observations (N = 177) than our false positive data set (N = 81) we conducted a bootstrapping analysis to investigate if any effect present for true positive could be present in the false positive model but remain undetected due to differences in sample size. We generated 500 bootstrap samples from the false positive logistic regression and refitted the model for every sample. We extracted the coefficient of the area effect for each sample and calculated the proportion of bootstrapped coefficients greater or equal to the observed area coefficient of the true positive model to assess the likelihood that a similar effect could be present in our false positive data (see supplementary material).

### Wave number and decision time

Furthermore, we explored differences between true and false positives through the influence of shoal size area on repeat wave numbers and decision time. First, we fitted a GLMM with a negative binomial error distribution and log link function, *response type* (true positive/false positive) as fixed effect and *wave number* as our response variable. To infer the influence of shoal size within the respective response type (true positive or false positive) we fitted two similar models on subsets containing only either true or false positive observations. To explore the influence of shoal size on decision time a second model was fitted with a gaussian error distribution (identity link) and *time to first repeat wave* as response variable. *Time to first repeat wave*, a proxy for the collective’s decision time, was log transformed and both models included *response type* (true positive/false positive) as additional fixed effect.

### Influence on predator performance

Lastly, we explored the effect of the focal shoal’s response behaviour on the predator behaviour. Models were therefore fitted on a subset with actual predator attacks only (i.e., true positives and false negatives). All models included sampling day and individual sequence as random effects. In a first model *predator success* (yes/no) was set as the response variable in a logistic regression and the *number of observed waves, shoal size area, decision time and attack in sequence* as fixed effects. In a second model *waiting time* (log transformed time in s) was set as response variable with *number of observed waves, success, decision time* and *shoal size area* as fixed effects.

All models were fitted with and without an interaction between main fixed effects of interest and shoal size area. Unless specified otherwise non-significant interaction effects were stepwise removed starting with the least significant interaction. Inference of the final models was assessed using the DHARMa library ^56^ and models were considered fitting if the KS, dispersion and outlier tests were not significant, and if no patterns were observable in the residual vs. prediction plots. All models were checked for overdispersion, temporal autocorrelation and zero inflation (negative binomial models only) with the respective build-in DHARMa functions. Visualisation of data was produced with ggplot2.

## Acknowledgement

We would like to thank the director at the CIIEA Centro de Investigación e Innovación para la Enseñanza y el Aprendizaje, Teapa, Tabasco field station for hosting us. We further thank Ildiko Rolfes and Sara Günter for helping in data collection and Charlotte Steven and Anna Helmke for assistance during the fieldwork.

## Author contributions

J.K., P.R. D.B. and L. A-R. acquired funding. K.P., J.K., D.B. designed the field study. K.P., J.K., A. J-L., J. E. J-J., Y. S., and C.V. conducted the fieldwork. K.P., J.K, Y.S. and D.B. outlined the data analysis. K.P. and D.B. conducted the data analysis. K.P. wrote the first manuscript draft. All authors commented on the manuscript and substantially contributed to the final version.

## Data and Code availability

Raw data for behavioural analysis of decision performance is supplied with the supplementary material. Reproducible code and data for density analysis will be made available by the corresponding author upon reasonable request.

## Ethical statement

Field observations reported in this study were carried out in accordance with the recommendations of ‘Guidelines for the treatment of animals in behavioural research and teaching’ (published in Animal Behaviour 1997) and under the authorization of the Mexican government (H. Ayuntamiento Constitucional Tacotalpa, Direction Fomento Economico y Turismo; DGOPA.09004.041111.3088, PRMN/DGOPA-003/2014, PRMN/DGOPA-009/2015, and PRMN/DGOPA-012/2017, PRMN/DGOPA-2018 issued by SAGARPACONAPESCA-DGOPA).

