## Supplementary material for "Better and faster collective decisions by larger fish shoals in the wild": SI Material

**Supplementary Information**

*Ground truthing of rectification and measurements*

As our approach relies heavily on measurements obtained from rectified videos, we applied a multilevel ground truthing to account for sufficient accuracy in our measurements of shoal size and bird speed. In a first step we acquired video recordings in which flat objects of known size were filmed in different positions and then repeatedly measured for length and area in rectified videos (see Fig S1a). In a second step we measured marked distances at our field sites at water level and remeasured the same distances in rectified videos recorded from two fundamentally different angles at the river shoreline (see FigS1b; c). Measurements of objects of known size had a mean measurement error of – 0.124 ± 0.11 m (N = 5) for length measurements and a mean measurement error of 0.0016 ± 0.013 m^2^ for area measurements (N = 10). Measurements of known distances in the field from different perspectives revealed a mean measurement error of -0.08 ± 0.27 m (N = 28 measurements).

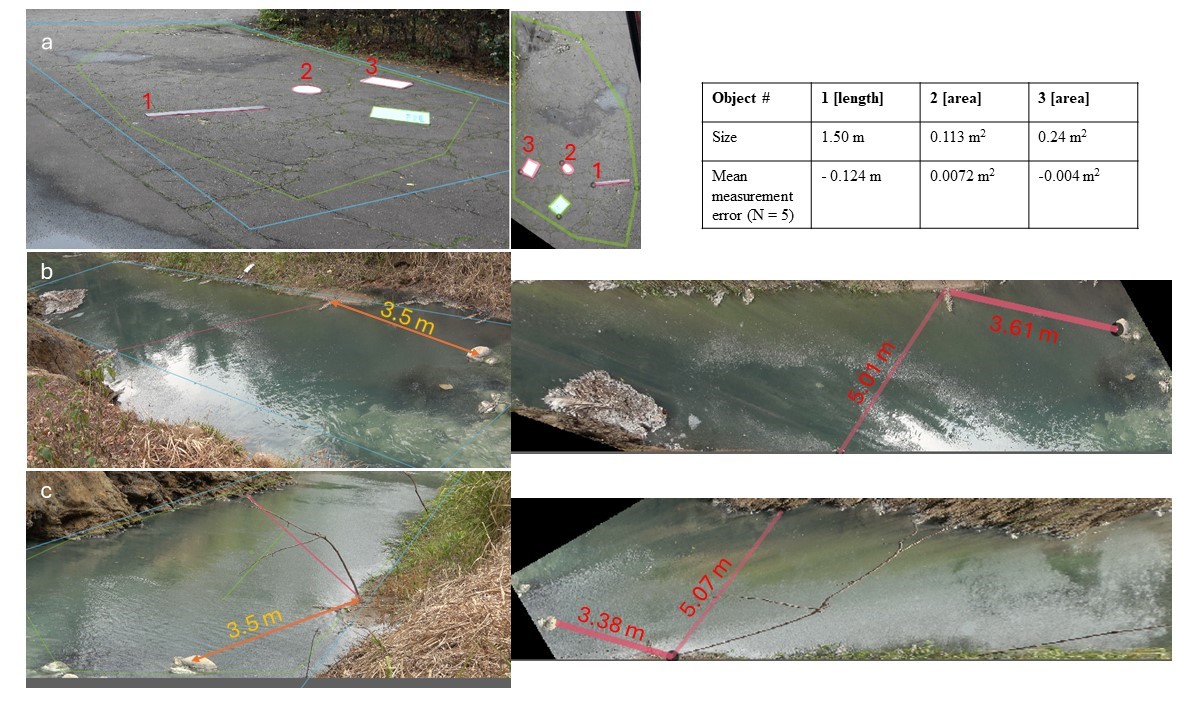

**Figure S1**: Ground truthing of distance measurements. In a first steps repeated measurements of length and area of objects of known size were performed in rectified video recordings (a). In a second step we measured distances at our field sites (example: b,c: orange arrow in left panels) and measured the same distance in rectified videos (b,c: right panels) filmed from different perspectives (b,c left panels).

*Field sites*

Table S1: Location of field sites at which behavioural data was gathered.

| Site | Location (lon,lat) | N _Attacks_ | N _Overflights_ | N _Recording days_ |
| --- | --- | --- | --- | --- |
| 1 | 17.557412, -93.006040 | 41 | 22 | 2 |
| 2 | 17.552779, -92.997249 | 99 | 52 | 3 |
| 3 | 17.554398, -93.004935 | 37 | 7 | 1 |

*Kingfisher attacks as control group*

We randomly selected 47 kingfisher attacks from one sampling day per location to compare the true positive rate as function of group size between their conspicuous plunge diving attacks (See figure S2 below) and the great kiskadees ambiguous overflight attacks (See Figure 2 main text). We included attacks of two kingfisher species: The amazon kingfisher (*Chloroceryle amazonae; N = 10*) and the green kingfisher (*Chloroceryle americana; N = 37*). As our field recordings consist of complete days, kingfisher hunting behaviour is occasionally present on our video footage but used as control group representing a multimodal predatory cue in this study. For kingfisher attacks we recorded the number of repeat waves, shoal size area and if the response by the prey represented a true positive or false negative. The true positive rate for kingfishers was found to be 100% accurate with all attacks classified as true positives. A kingfisher attack produced on average 31.8 ± 22.3 waves, with a maximum of 91 waves and a minimum of 2.

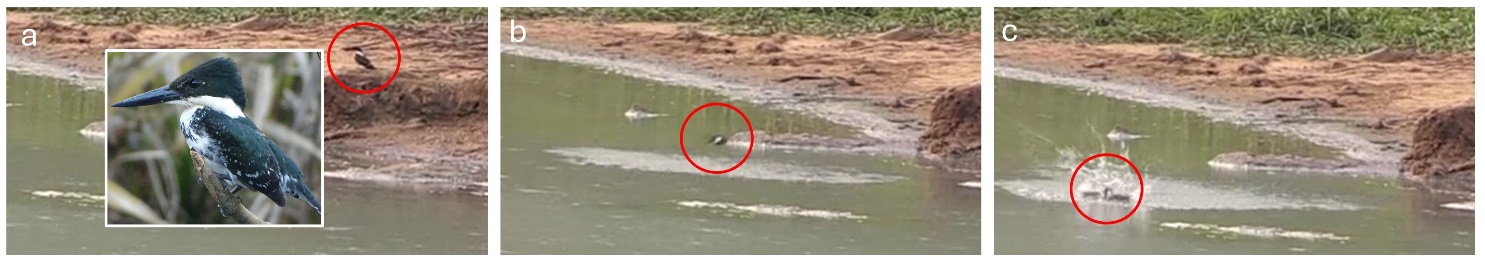

Figure S2: Plunge diving attack by green kingfisher. During a typical attack kingfishers (red circle) attack from elevated perches (a) and enter the water in full flight (b, c). This produces a multimodal cue that is easily detected as an attack by prey shoals.

*Bootstrapping simulation*

To account for differences in sample size between attacks and flybys we performed a bootstrapping simulation in which the false positive logistic regression was refitted 500 times and compared the distribution of coefficients to the observed coefficient of the true positive model. Notably, only 0.4% of the bootstrapped samples produced area coefficients that were greater than or equal to the observed true positive coefficient (See Fig S2).

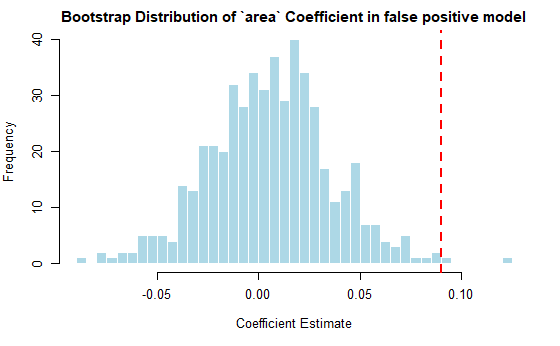

**Figure S3:** Bootstrap distribution of 500 area coefficients from the false positive logistic regression. The red dashed line represents the observed area coefficient (0.09) from the true positive model (N = 177). The histogram shows that only 0.4% of the bootstrapped coefficients are greater than or equal to the observed area coefficient for true positive rate, indicating that such an effect is unlikely to be observed in the false positive data.

*Relationship of shoal size area and density*

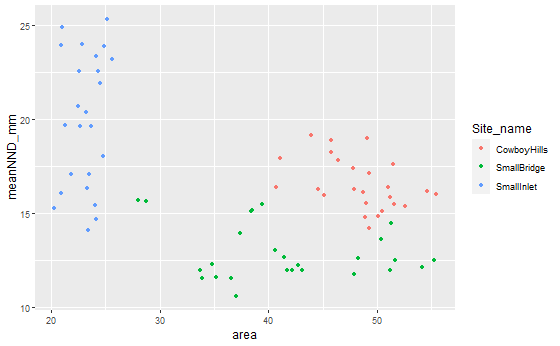

**Figure S4:** Relationship of shoal size area and fish density expressed as nearest neighbour distance (NND). Dots represent mean NND´s measured between all annotated fish in a still frame from high detail video recordings and the measured shoal size from wide angle recordings at the same time. Colours represent our three field sites. Mean NND´s differed significantly between field sites. There was no relationship between mean NND and area.

*Attack and flyby speeds*

To compare speeds between flybys and attacks by the great kiskadee, as well as between flybys by the great kiskadee and other birds we measured their flight distance per time unit in rectified video recordings. For attacks this was done with the point at which the birds beak enters the water surface as the central event. From there we measured the bird’s distance as a straight line on the water surface ten frames before (pre-attack speed) and after (post-attack speed) the point of attack. For flybys we started measuring the flight distance the same way as soon as the bird crossed the shoal outline. We measured 16 pre-attack and 15 post-attack speeds by the great kiskadee as well as 10 flybys by the kiskadee and 7 by other birds. We checked for differences between general attack and flyby speed, pre-attack and post-attack speed and inter-species flyby speed by applying Welch´s t-test. Flybys were overall significantly faster with a mean speed of 14.1 ms-1 than the average attack with 10.9 ms^-1^ (Welch-t (-4.37): 21.635, p < 0.001). Flybys by the kiskadee were not significantly different in speed from other birds (Welch-t (-1.23): 7.22, p = 0.26) and neither were kiskadee pre-attack and post-attack flight speeds (Welch-t (0.50): 28.89, p = 0.61).

*Model details*

|  | **M1 Density (meanNND_mm)** | | |
| --- | --- | --- | --- |
| *Predictors* | *Estimates* | *CI* | *p* |
| (Intercept) | 19.15 | 14.03 – 24.28 | **<0.001** |
| area | -0.05 | -0.16 – 0.05 | 0.300 |
| Site name [SmallBridge] | -3.96 | -5.39 – -2.53 | **<0.001** |
| Site name [SmallInlet] | 2.09 | -0.84 – 5.02 | 0.161 |
| **Random Effects** | | | |
| σ^2^ | 5.16 | | |
| τ_00_ _Site_name:day_ | 0.00 | | |
| N _Site_name_ | 3 | | |
| N _day_ | 6 | | |
| Observations | 76 | | |
| Marginal R^2^ / Conditional R^2^ | 0.620 / NA | | |

|  | **M2: Polarisation (meanPolarization_total)** | | |
| --- | --- | --- | --- |
| *Predictors* | *Estimates* | *CI* | *p* |
| (Intercept) | 0.30 | -0.74 – 1.35 | 0.572 |
| area | -0.00 | -0.02 – 0.02 | 0.755 |
| Site name [SmallBridge] | -0.28 | -1.32 – 0.76 | 0.599 |
| Site name [SmallInlet] | 1.52 | -1.24 – 4.28 | 0.280 |
| area Ã— Site name [SmallBridge] | 0.02 | -0.00 – 0.04 | 0.121 |
| area Ã— Site name [SmallInlet] | -0.06 | -0.19 – 0.06 | 0.336 |
| **Random Effects** | | | |
| σ^2^ | 0.02 | | |
| τ_00_ _Site_name:day_ | 0.00 | | |
| ICC | 0.10 | | |
| N _Site_name_ | 3 | | |
| N _day_ | 6 | | |
| Observations | 76 | | |
| Marginal R^2^ / Conditional R^2^ | 0.656 / 0.691 | | |

|  | **M3: true_positive** | | |
| --- | --- | --- | --- |
| *Predictors* | *Odds Ratios* | *CI* | *p* |
| (Intercept) | 0.06 | 0.01 – 0.50 | **0.009** |
| area | 1.09 | 1.05 – 1.14 | **<0.001** |
| location [SmallBridge] | 0.17 | 0.06 – 0.51 | **0.002** |
| location [SmallInlet] | 1.97 | 0.56 – 6.93 | 0.290 |
| attack in bout | 0.97 | 0.85 – 1.09 | 0.567 |
| **Random Effects** | | | |
| σ^2^ | 3.29 | | |
| τ_00_ _location:day_ | 0.00 | | |
| N _location_ | 3 | | |
| N _day_ | 6 | | |
| Observations | 177 | | |
| Marginal R^2^ / Conditional R^2^ | 0.226 / NA | | |

|  | **M4: false_positive** | | |
| --- | --- | --- | --- |
| *Predictors* | *Odds Ratios* | *CI* | *p* |
| (Intercept) | 0.31 | 0.02 – 5.19 | 0.419 |
| area | 1.00 | 0.95 – 1.06 | 0.863 |
| location [SmallBridge] | 0.85 | 0.26 – 2.82 | 0.787 |
| location [SmallInlet] | 1.87 | 0.28 – 12.29 | 0.516 |
| species2 [other] | 1.64 | 0.61 – 4.40 | 0.323 |
| **Random Effects** | | | |
| σ^2^ | 3.29 | | |
| τ_00_ _location:day_ | 0.00 | | |
| N _location_ | 3 | | |
| N _day_ | 6 | | |
| Observations | 81 | | |
| Marginal R^2^ / Conditional R^2^ | 0.023 / NA | | |

|  | **M5: repeat wave number (re_wav)** | | |
| --- | --- | --- | --- |
| *Predictors* | *Incidence Rate Ratios* | *CI* | *p* |
| (Intercept) | 0.19 | 0.04 – 0.94 | **0.042** |
| area | 1.04 | 1.01 – 1.07 | **0.005** |
| group [True Positives] | 3.01 | 1.68 – 5.41 | **<0.001** |
| location [SmallBridge] | 0.55 | 0.27 – 1.13 | 0.103 |
| location [SmallInlet] | 2.22 | 0.90 – 5.45 | 0.083 |
| **Random Effects** | | | |
| σ^2^ | 0.78 | | |
| τ_00_ _location:day_ | 0.09 | | |
| ICC | 0.10 | | |
| N _location_ | 3 | | |
| N _day_ | 6 | | |
| Observations | 153 | | |
| Marginal R^2^ / Conditional R^2^ | 0.315 / 0.386 | | |

|  | **M5.1: repeat wave number for true positive (re_wav_TP)** | | |
| --- | --- | --- | --- |
| *Predictors* | *Incidence Rate Ratios* | *CI* | *p* |
| (Intercept) | 0.42 | 0.07 – 2.31 | 0.315 |
| area | 1.05 | 1.01 – 1.08 | **0.003** |
| location [SmallBridge] | 0.48 | 0.21 – 1.08 | 0.077 |
| location [SmallInlet] | 2.43 | 0.88 – 6.68 | 0.085 |
| **Random Effects** | | | |
| σ^2^ | 0.73 | | |
| τ_00_ _location:day_ | 0.12 | | |
| ICC | 0.14 | | |
| N _location_ | 3 | | |
| N _day_ | 6 | | |
| Observations | 127 | | |
| Marginal R^2^ / Conditional R^2^ | 0.202 / 0.315 | | |

|  | **M5.2: repeat wave number for false positive (re_wav_FP)** | | |
| --- | --- | --- | --- |
| *Predictors* | *Incidence Rate Ratios* | *CI* | *p* |
| (Intercept) | 0.36 | 0.02 – 8.38 | 0.523 |
| area | 1.01 | 0.95 – 1.07 | 0.692 |
| location [SmallBridge] | 1.81 | 0.42 – 7.79 | 0.423 |
| location [SmallInlet] | 0.88 | 0.08 – 9.17 | 0.912 |
| **Random Effects** | | | |
| σ^2^ | 1.11 | | |
| τ_00_ _location:day_ | 0.00 | | |
| N _location_ | 3 | | |
| N _day_ | 6 | | |
| Observations | 26 | | |
| Marginal R^2^ / Conditional R^2^ | 0.136 / NA | | |

|  | **M6: Decision Speed (log(t_first))** | | |
| --- | --- | --- | --- |
| *Predictors* | *Estimates* | *CI* | *p* |
| (Intercept) | 5.44 | 4.99 – 5.90 | **<0.001** |
| area | -0.01 | -0.02 – -0.00 | **0.002** |
| location [SmallBridge] | 0.19 | 0.01 – 0.37 | **0.042** |
| location [SmallInlet] | -0.07 | -0.30 – 0.16 | 0.530 |
| group [True Positives] | -0.11 | -0.31 – 0.08 | 0.255 |
| **Random Effects** | | | |
| σ^2^ | 0.19 | | |
| τ_00_ _location:day_ | 0.00 | | |
| N _location_ | 3 | | |
| N _day_ | 6 | | |
| Observations | 148 | | |
| Marginal R^2^ / Conditional R^2^ | 0.085 / NA | | |

|  | **M7: Predator success (succ_bin)** | | |
| --- | --- | --- | --- |
| *Predictors* | *Odds Ratios* | *CI* | *p* |
| (Intercept) | 0.94 | 0.01 – 167.07 | 0.980 |
| wave before | 1.00 | 0.90 – 1.10 | 0.940 |
| attack in bout | 0.95 | 0.79 – 1.15 | 0.613 |
| location [SmallBridge] | 0.73 | 0.16 – 3.29 | 0.678 |
| location [SmallInlet] | 0.76 | 0.09 – 6.13 | 0.794 |
| t first | 1.00 | 0.99 – 1.01 | 0.783 |
| area | 0.99 | 0.91 – 1.09 | 0.884 |
| **Random Effects** | | | |
| σ^2^ | 3.29 | | |
| τ_00_ _location:day_ | 0.09 | | |
| τ_00_ _unique_bout_ | 0.00 | | |
| N _location_ | 3 | | |
| N _day_ | 6 | | |
| N _unique_bout_ | 31 | | |
| Observations | 85 | | |
| Marginal R^2^ / Conditional R^2^ | 0.016 / NA | | |

|  | **M8: Predator waiting time (log(time_lag))** | | |
| --- | --- | --- | --- |
| *Predictors* | *Estimates* | *CI* | *p* |
| (Intercept) | 4.92 | 3.29 – 6.56 | **<0.001** |
| wave before | 0.04 | -0.00 – 0.07 | 0.071 |
| t first | 0.00 | -0.00 – 0.00 | 0.170 |
| location [SmallBridge] | -0.44 | -1.06 – 0.18 | 0.167 |
| location [SmallInlet] | -0.76 | -1.53 – 0.01 | 0.054 |
| area | -0.02 | -0.05 – 0.01 | 0.202 |
| **Random Effects** | | | |
| σ^2^ | 0.76 | | |
| τ_00_ _location:day_ | 0.00 | | |
| τ_00_ _unique_bout_ | 0.12 | | |
| N _location_ | 3 | | |
| N _day_ | 6 | | |
| N _unique_bout_ | 31 | | |
| Observations | 91 | | |
| Marginal R^2^ / Conditional R^2^ | 0.174 / NA | | |
